# Improving fMRI-based Autism Spectrum Disorder Classification with Random Walks-informed Feature Extraction and Selection

**DOI:** 10.1101/2023.07.05.547843

**Authors:** Roberto C. Sotero, Jose M. Sanchez-bornot, Yasser Iturria-medina

## Abstract

Functional magnetic resonance imaging (fMRI) is a non-invasive technique measuring brain activity by detecting blood flow changes, enabling the study of cognitive processes and brain states. However, the high dimensionality of resting-state (rs) fMRI data poses challenges for machine learning applications. Feature extraction (FE) and feature selection (FS) are critical for developing efficient machine learning models. Transforming raw data into meaningful features and selecting the most relevant ones, allows models to achieve improved generalization, accuracy, and robustness. Previous studies demonstrated the effectiveness of FE and FS methods for analyzing rs-fMRI data for Autism Spectrum Disorder (ASD) classification. In this study, we apply a random walks technique for correlation-based brain networks to extract features from rs-fMRI data, specifically the number of random walkers on each brain area. We then select significant features, i.e., brain areas with a statistically significant difference in the number of random walkers between neurotypical and ASD subjects. Our random walks-based FE and FS approach reduces the number of brain areas used in the classification and converts the functional connectivity matrix into a manageable vector, enabling faster computation. We examined 16 pipelines and tested support vector machines (SVM) and logistic regression for classification, identifying the optimal pipeline to consist of no filtering, no global signal regression (GSR), and FS, achieving a 76.54% classification accuracy with SVM. Our findings suggest that random walks capture a wide range of interactions and dynamics in brain networks, providing a deeper characterization of their structure and function, ultimately enhancing classification performance.

**CCS CONCEPTS:** Computing methodologies→Machine learning

## 1 INTRODUCTION

Resting-state functional magnetic resonance imaging (rs-fMRI) data are 4-dimensional (4D) volumes (3 spatial dimensions + 1 temporal dimension) that serve as an indirect measure of neuronal activity [1], [2], and have been utilized to investigate information flow between brain networks and regions in both healthy and diseased states [3]. However, the high dimensionality of rs-fMRI data makes it challenging to directly analyse for machine learning applications. To address this issue, most studies subject the 4D fMRI data to a transformation process that encompasses preprocessing steps like slice timing correction, motion correction, smoothing, temporal filtering, global signal regression (GSR), brain parcellation, and extraction of the mean time series from each region of interest (ROI) [4], [5]. The resulting time series can then be used to examine functional connectivity between different brain areas [6]. Nonetheless, depending on the chosen brain parcellation and the number of recorded time points, the transformed data may still be too large for analysis. Additionally, not all time series derived from various brain regions may offer valuable information when used for tasks such as classifying specific disorders, highlighting the need for careful consideration when selecting brain regions and time series for analysis in machine learning experiments [7].

Feature extraction (FE) and feature selection (FS) are two crucial steps in developing efficient and effective machine learning models [8]. FE involves transforming or generating new features from raw data to better represent underlying patterns, thereby facilitating learning algorithms in capturing relationships between inputs and outputs [9]. Conversely, FS aims to identify and retain the most pertinent and informative features from the extracted or transformed set, ultimately reducing noise, enhancing model interpretability, and decreasing computational complexity [10]. Integrating FE and FS allows for a more streamlined, optimized learning process that improves model performance. By meticulously extracting meaningful features and selecting only the most relevant among them, machine learning models can attain improved generalization, accuracy, and robustness across various complex tasks, leading to more dependable and efficient solutions [11].

Several studies have showcased the efficacy of FE and FS methods in the analysis of rs-fMRI data for Autism Spectrum Disorder (ASD) classification, specifically utilizing the ABIDE dataset [12]. For instance, Zhang et al. [13] employed an F-score FS technique and achieved a 70.9% accuracy on a sample of 1112 subjects. Shao et al. [14] integrated a deep FS network with a graph convolutional network, resulting in a 79.5% accuracy for 871 subjects. Another recent study [15] used support vector machines (SVM) for both classification and FS, attaining 96.15% accuracy on a smaller subset of 84 subjects. Additionally, another research [16] implemented a FS approach based on multiple autoencoders, achieving 86.36% accuracy for 110 subjects. Kazeminejad and Sotero [17] used graph theory metrics for FE obtaining 95.12% accuracy when partitioning the data into five age groups. These studies underscore the potential of employing FE and FS techniques for ASD classification using the ABIDE dataset.

In this study, we apply a recently introduced random walks technique for correlation-based brain networks [18], to extract features from rs-fMRI data, specifically, the number of random walkers on each brain area which may be an indication of the role of the region as a hub for brain communication. We then select the most salient regional features, i.e., brain areas with a statistically significant difference in the number of random walkers between neurotypical and ASD subjects. The random walk-based FE and FS approach proposed here reduces the number of brain areas used in the classification to less than 50 areas, from the original 116 in the anatomical automatic labeling (AAL) atlas [19]. This method also dramatically reduces the number of functional connectivity features, enabling faster computation of the results. We examined 16 different pipelines, comprising all combinations of bandpass filtering (0.01 - 0.1Hz) and without filtering, with and without GSR, and with (all features, features where ASD>neurotypical, features where ASD<neurotypical) and without FS. For classification, we tested SVM and logistic regression, finding the optimal pipeline to consist of no filtering, no GSR, and FS, achieving a classification accuracy of 76.54 % with SVM. Our findings suggest that random walks capture a broad spectrum of interactions and dynamics within the brain network, offering a more comprehensive understanding of its structure and function. This insight informs the classification method, improving classification accuracies.

## 2 DATA PREPROCESSING

This study used rs-fMRI data from 948 subjects (ASD: 457, neurotypical: 491) obtained from the ABIDE Preprocessed Initiative [12]. To ensure that our results were not affected by any custom preprocessing pipeline, we used the preprocessed data provided by ABIDE in the C-PAC pipeline [20]. The preprocessing included the following steps. The Analysis of Functional Neuro Images (AFNI) software [21] was used for removing the skull from the images. The brain was segmented into three tissues using FMRIB Software Library (FSL) [22]. The images were then normalized to the MNI 152 stereotactic space [23], [24] using Advanced Normalization Tools (ANTs) [25]. Functional preprocessing included motion and slice-timing correction and the normalization of voxel intensity. Nuisance signal regression included 24 parameters for head motion, CompCor [26] with five principal components for tissue signal in Cerebrospinal fluid and white matter, linear and quadratic trends for Low-frequency drifts. These images were then co-registered with their anatomical counterpart by FSL. They were then normalized to the MNI 152 space using ANTs. The average voxel activity in each Region of Interest (ROI) of the AAL atlas [19] was then extracted as the time series for that region.

Due to the controversies surrounding the use of bandpass filtering [27] and GSR [28], we considered different preprocessing strategies: either with bandpass filtering (0.01-0.1Hz) or without bandpass filtering, and with or without GSR. Different ABIDE sites used different acquisition protocols resulting in rs-fMRI data with different sampling rates [29]. Thus, we resampled each time series using a resolution of 2 s to have them all in the same time scale. We then applied a Z-score normalization (subtracted the mean and divided the result by the standard deviation) to each time series. Training neural networks frequently requires data augmentation to avoid overfitting. As different subjects had varying time series lengths depending on the originating ABIDE site [29], we opted to segment all time-series into consecutive sequences of a fixed length (*T* = 2 min), thereby augmenting the number of inputs for each subject. Functional connectivity matrices were subsequently generated by computing the Pearson correlation coefficient between every pair of time series. Non-significant connections were set to zero using a False Discovery Rate (FDR)-adjusted p-value of 0.05.

## 3 FEATURE EXTRACTION AND SELECTION THROUGH RANDOM WALKS

We use a random walks model to examine the functional connectivity network, aiming to obtain fewer, yet more effective features for classification. Given that our functional connectivity networks rely on correlation, we apply a recently proposed routing strategy that considers the sign of the connection [18]. In this random walk model the transition probabilities are calculated based on the weights in the functional connectivity matrix (*C*) and a similarity measure between the activities of the source and destination nodes. The primary concept is to promote transitions between nodes with similar activities while considering the positive or negative weights in the network. We define the activity in each node as the number of walkers visiting the node. At the beginning (t = 0), we use a uniform random distribution to assign each of the *K* walkers to one of the 116 brain areas, resulting in an initial activity *A*_*i*_(0) for each node *i*. Given the activities *A*_*i*_(0) the transition probabilities *P*_*ij*_ (*t* + 1) and the next activities *A*_*i*_(*t* + 1) are computed as follows:

1. For each node *i*, we calculate the adjusted weights for all its neighboring nodes j as follows:
  a. We introduce the similarity measure *S*_*ij*_ (*t*) between the activity of node *i* and node *j* at time *t* using the formula:

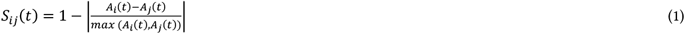

To normalize the difference between the activities of the nodes, we divide by the maximum of the activities *max* (*A*_*i*_(*t*), *A*_*j*_ (*t*)). The similarity *S*_*ij*_ (*t*) ranges between 0 and 1. A value closer to 1 indicates that the activities of nodes *i* and node *j* are more similar, while a value closer to 0 indicates that the activities are more dissimilar.
  b. If the weight *C*_*ij*_ is positive, the adjusted weight is 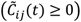:

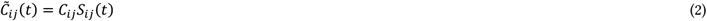

If the weight *C*_*ij*_ is negative, the adjusted weight is:

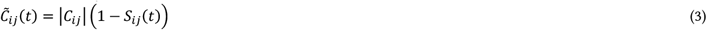
3. The transition probability *P*_*ij*_ (*t* + 1) of a walker moving from node *i* to each neighboring node *j* in the next time step *t* + 1 is computed as:

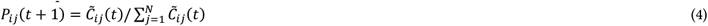

Thus, a walker is more likely to travel through a positive connection *C*_*ij*_ if the activities in nodes *i* and *j* are similar, while it is more likely to take a negative connection if the activities are dissimilar.
4. The random walkers move based on the computed probabilities *P*_*ij*_ (*t* + 1), and the next activites *A*_*i*_(*t* + 1) are the new number of walkers at each node.

Figure 1 illustrates the FE method for a single subject. Figure 1A presents the rs-fMRI signals for the 116 brain areas, while Figure 1B displays the functional connectivity matrix calculated using the Pearson correlation between all pairs of time-series in Figure 1A. Next, we placed 10_5_ random walkers on the network, allowing them to explore it for 1100 time steps according to the rules in equations (1)-(4), and recorded the number of walkers in each node, i.e, the activity, *A*_*i*_(*t* + 1) (Figure 1C). The first 100 points were discarded to eliminate transient behaviour. Figure 1D shows the average activity across all time steps, which is a vector of 116 values that will serve as the new features.

**Figure 1:**
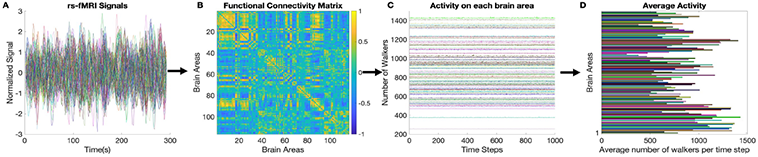
Feature extraction method for a single subject. (a) rs-fMRI signals for 116 brain areas. (b) Functional connectivity matrix calculated using Pearson correlation between all pairs of time series from (a). (c) Number of walkers in each node after placing 10^5^ random walkers and letting them move for 1100 time steps. The initial 100 values were discarded. (d) Average activity across all time steps, represented as a vector of 116 values, which serve as the new features.

After calculating the average activity in each brain area for all ASD and neurotypical subjects, we determined the activity difference between the two groups for each brain area and performed a t-test to identify areas with significant differences (p<0.05). Figure 2A and 2B display the number of walkers averaged across subjects and time steps for neurotypical and ASD conditions, respectively, ordered in descending order. ‘L’ and ‘R’ represent the left and right brain hemispheres, respectively. As shown in the figure, the walkers distribute according to different patterns for each condition. Although ‘Cerebellum 6 L’ area had the highest number of walkers in both cases, the other areas generally differ. Figure 2C presents only the 43 areas with significant differences between ASD and neurotypical conditions. These are the features ‘selected’ by the random walkers. The two brain areas with the largest differences where ‘Occipital Inf R’ (Autism>Neurotypical) and ‘Heschl R’ (Autism<Neurotypical). Both the Inferior Occipital and Heschl’s gyrus areas have been reported to show alterations in individuals with ASD. The Inferior Occipital is involved in visual processing, and individuals with ASD have been found to exhibit atypical visual processing and functional connectivity within this region [30]. Additionally, the Heschl’s area which is associated with auditory processing, has also been reported to have connectivity differences in individuals with autism [31], potentially contributing to the sensory processing abnormalities often observed in ASD.

**Figure 2:**
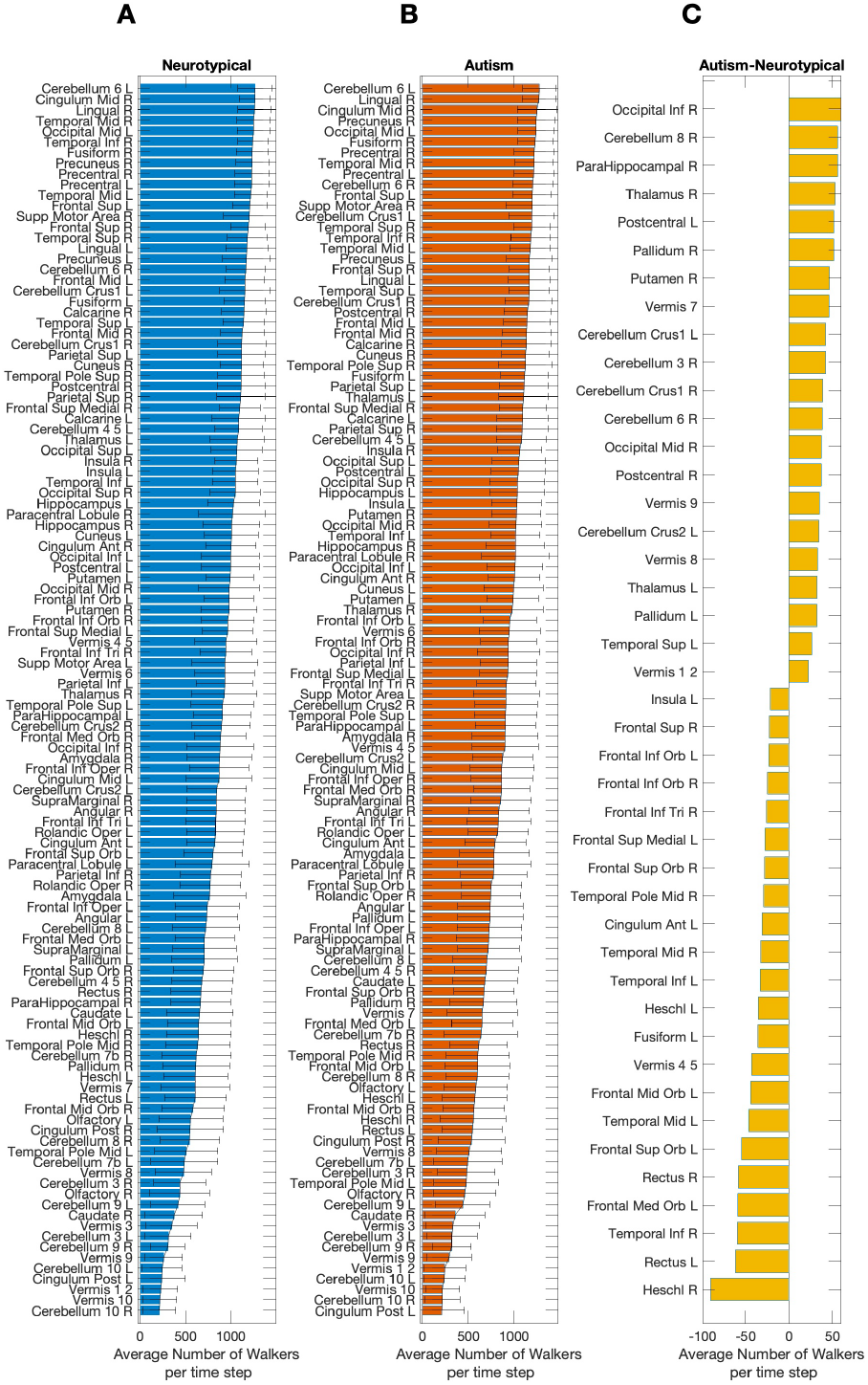
Feature selection methodology. (a) Number of walkers in each of the 116 brain areas of the AAL atlas averaged across neurotypical subjects and time steps, and ordered in descending order. (b) Number of walkers averaged across ASD subjects and time steps, sorted in descending order. ‘L’ and ‘R’ denote left and right hemispheres, respectively. (6) Brain areas with statistically significant activity differences between the two groups, which are the features ‘selected’ by the random walkers.

## 4 TRAINING AND CLASSIFICATION RESULTS

### 4.1 Hyperparameter selection

We trained a linear model using stochastic gradient descent (SGD) for each of the different pipelines in MATLAB R2023a (The MathWorks Inc., Natick, MA, USA) with the *fitclinear* function. We selected two hyperparameters: the model type (either SVM or logistic regression) and the regularization parameter lambda (*λ* ranges from 10^−5^ to 10^5^) that controls the trade-off between fitting the data well and keeping the model simple. Smaller values of *λ* allow more complex models and larger values promote simpler models. To tune the hyper-parameters, we used Bayesian optimization with the expected improvement acquisition function [32].

### 4.2 Feature selection and classification within a nested cross-validation framework

First, the preprocessed fMRI dataset from ABIDE is transformed into new features (i.e, the FE step) using the signed random walk [18] described in equations (1)-(4) and illustrated in Figure 1. To evaluate the different pipelines, we employ a nested cross-validation (CV) approach. Nested CV involves a double loop (Figure 3): an outer loop to assess the quality of the model and an inner loop used for model/parameter selection. We repeat the outer loop ten times, generating ten distinct test sets. Additionally, we partition the outer train set into ten folds (inner loop) for each iteration. With 10 outer folds and 10 inner folds (Figure 3), each pipelines trains a total of 100 models. As we are using an augmented dataset, we ensure that all data belonging to the same subject appear together in either training, validation, or testing sets. Importantly, to prevent data leakage, we compute statistically significant differences in activities between ASD and neurotypical subjects using only the random walkers’ activities from the training set, and then select the features (i.e, brain areas). We apply the selected brain areas to the testing set and subsequent steps, allowing the selected features to differ in each external loop.

**Figure 3:**
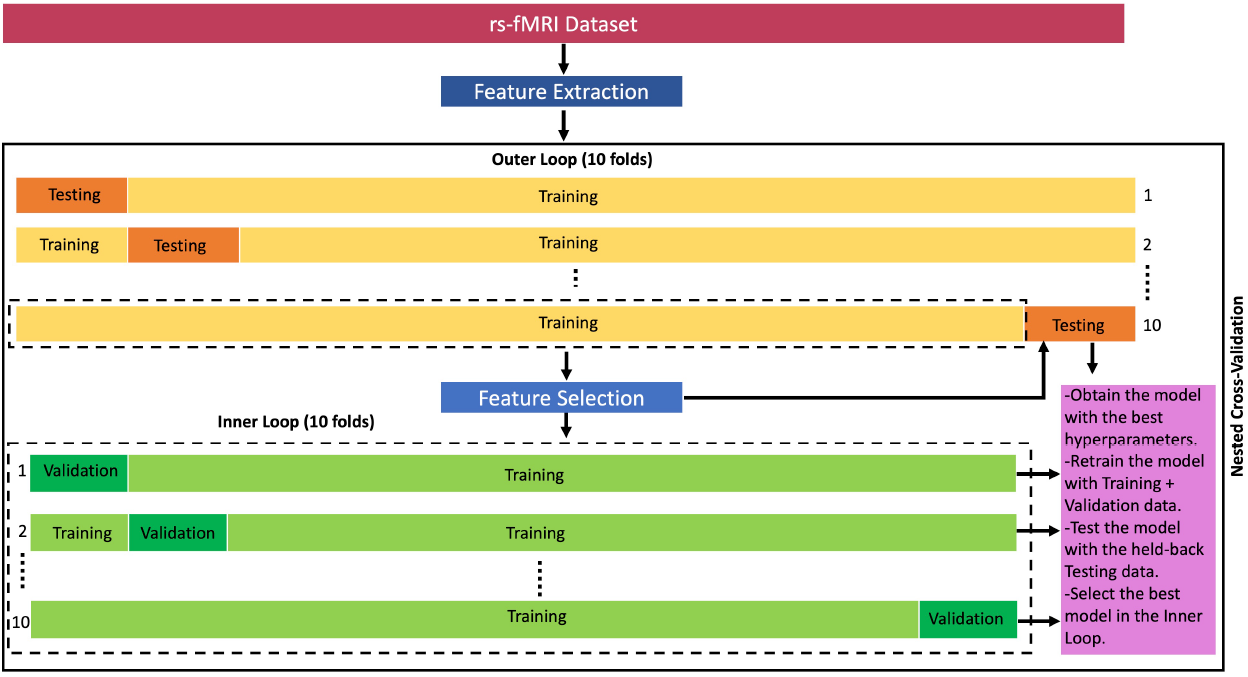
Nested cross-validation approach for model selection and validation. The approach consists of a double-loop scheme, with an outer loop assessing model quality and an inner loop for model/parameter selection. The outer loop is repeated ten times, generating ten different test sets, while the outer training set is split into ten folds (inner loop) for each iteration. To prevent data leakage, we select features independently for each training set. A total of 100 models are trained for each pipeline, and the best model is selected based on accuracy. The entire nested CV process is repeated 25 times and the results are averaged.

We go then to the inner loop, and for each fold, we obtain a model by tuning the hyper-parameters to minimize the validation loss and then retrain this model with all the data available in the loop (i.e., the training +validation data). We then compute the model accuracy (*ACC*) with the held-back testing data and select the model with the best accuracy in the inner loop. To calculate the accuracy we use 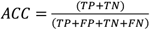, where TP is the number of true positives (the model correctly predicts ASD), TN is the number of true negatives (the model correctly predicts HC), FP is the number of false positives (the model incorrectly predicts ASD), and FN is the number of false negatives (the model incorrectly predicts HC). Accuracy is an overall measure of classification performance, with higher accuracy values being more valuable for practical medical diagnosis applications. Out of the 100 models in the nested CV, we select the best model. Finally, we repeat the nested CV 25 times and average the results.

### 4.3 Classification results

We trained models for 16 different pipelines, encompassing all combinations of bandpass filtering (0.01 - 0.1Hz) and without filtering, with and without GSR, and with (‘all’, ‘pos’, and ‘neg’) and without FS. The labels ‘all’, ‘pos’, and ‘neg’ in FS refer to the use of all significant features, of only the features where Autism>Neurotypical, and only the features where Autism<Neurotypical, respectively (see also Figure 2C. for an example).

Table 1 presents the classification results. The first column displays the pipeline label, the second column shows the average accuracy over the 25 repetitions of the nested CV scheme plus or minus the standard deviation. The third column indicates the best model out of the 25 repetitions, and the corresponding hyperparameters are provided in the fourth column. According to our results, the best model was ‘noGSR-noFilt-FS-all’ with a 76.54% accuracy, followed closely by ‘noGSR-Filt-FS-all’ with 76.21%. The table demonstrates that models using all significant features consistently performed better than those without FS. However, some models with the ‘pos’ case (like ‘GSR-noFilt-FS-pos’ with the best model having 73.21% accuracy), that resulted in a further decrease in the number of features, performed better than when using all features (‘GSR-noFilt-FS-all’ with the best model having 72.24%).

**Table 1:**
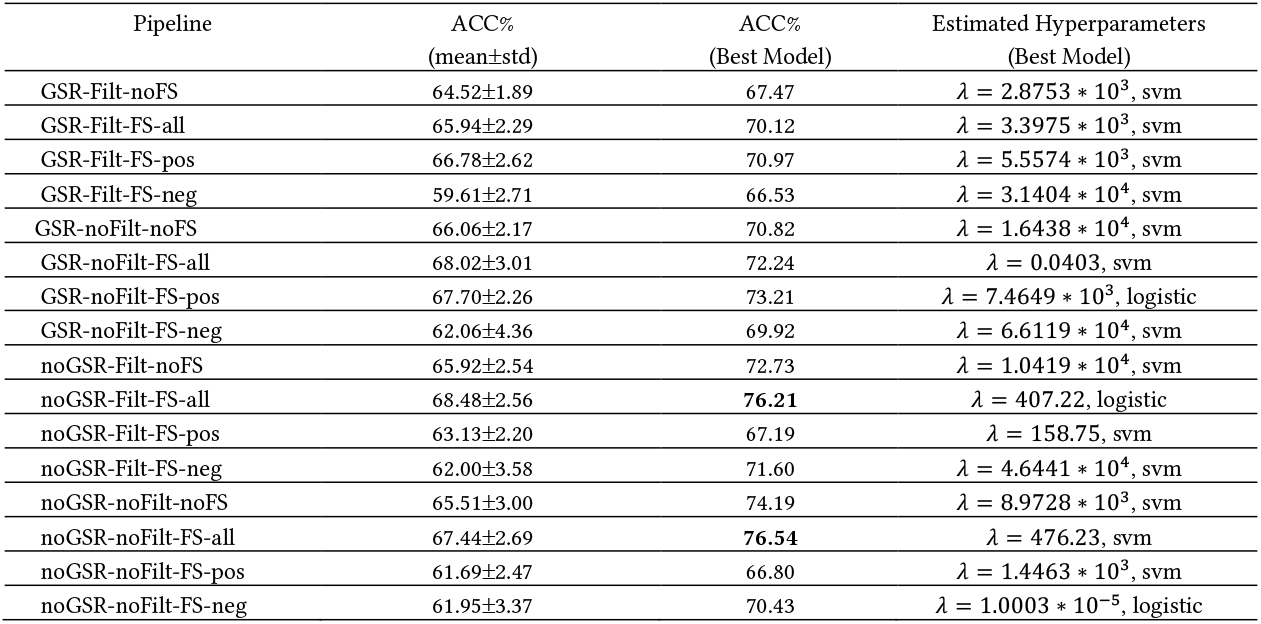
ASD classification results. The 16 preprocessing pipelines correspond to combinations of GSR, no GSR (noGSR), Filt (0.01-0.1Hz bandpass filtering), noFilt (not filtering), ‘all’ (all significant features), ‘pos’ (features where Autism>Neurotypical), and ‘neg’ (features where Autism<Neurotypical).

## 5 DISCUSSION

In this study, a linear model was employed for classifying ASD and neurotypical subjects based on rs-fMRI data. This approach, although insightful, had limitations such as the assumption of linearity, and the inability to capture feature interactions innately. Thus, there’s potential for enhancing performance by introducing nonlinear models like decision trees, random forests, or neural networks, capable of discerning complex data patterns. The application of deep learning methods, such as Convolutional Neural Networks, could also be explored [33]. Further feature engineering may reveal more informative features [34], but the increased complexity could render model interpretation more difficult. Future research should strive for a balance: enhancing accuracy while retaining model interpretability [35].

Moreover, to improve classification, we should consider limitations of the adaptive random walk model used in this study [18]. The use of static correlation-based functional connectivity matrices neglects the dynamics of brain activity [36]. Extending the random walks model to accommodate dynamic functional connectivity could provide more insightful understanding of time-varying brain network organization, enhancing classification performance.

Modelling studies have shown that GSR can introduce spurious anticorrelations not initially present in the modelled data [37]. However, proving the presence of spurious anticorrelations in actual data is challenging. The signed random walks method used here [18] can account for negative correlations. Our findings that models without GSR generally performed better than models with GSR is in contrast to another study [38] using graph theory and SVM, which found that including anticorrelations in their analysis improved ASD classification. This suggest that interpreting classification results from a highly heterogeneous dataset like ABIDE is largely dependent on the methods and features employed and should be approached with caution. Furthermore, to enhance classification performance and obtain reliable physiological information it is essential to address the presence of ASD subtypes within this dataset [39].

## 6 CONCLUSSIONS

In conclusion, this study presents a novel approach for enhancing ASD classification by combining signed random walks-informed feature extraction and selection with rs-fMRI data. Our findings demonstrate that the use of all significant features consistently yielded better results than not using FS. In some cases, models with only positive features (Autism>Neurotypical) performed better than those using all features, indicating the potential for further reducing the number of features required for classification. Further research could focus on exploring more advanced machine learning techniques and architectures and refining the random walks model. Moreover, investigating the impact of data heterogeneity and addressing ASD subtypes within the dataset could provide valuable insights into the development of more accurate and robust ASD classification models.

## ACKNOWLEDGMENTS

This work was supported by the Alberta Innovates LevMax program, Grant 222300868.

## REFERENCES

[1] B. Biswal, F. Z. Yetkin, V. M. Haughton, and J. S. Hyde, “Functional connectivity in the motor cortex of resting human brain using echo-planar MRI,” Magn Reson Med, vol. 34, no. 4, pp. 537–541, Oct. 1995, doi: 10.1002/mrm.1910340409.

[2] M. D. Fox and M. E. Raichle, “Spontaneous fluctuations in brain activity observed with functional magnetic resonance imaging,” Nat Rev Neurosci, vol. 8, no. 9, Art. no. 9, Sep. 2007, doi: 10.1038/nrn2201.

[3] R. C. Sotero, L. M. Sanchez-Rodriguez, M. Dousty, Y. Iturria-Medina, and J. M. Sanchez-Bornot, “Cross-frequency interactions during information flow in complex brain networks are facilitated by scale-free properties,” Frontiers in Physics, vol. 7, p. 107, 2019.

[4] R. C. Craddock, G. A. James, P. E. Holtzheimer, X. P. Hu, and H. S. Mayberg, “A whole brain fMRI atlas generated via spatially constrained spectral clustering,” Hum Brain Mapp, vol. 33, no. 8, pp. 1914–1928, Aug. 2012, doi: 10.1002/hbm.21333.

[5] J. D. Power et al., “Functional network organization of the human brain,” Neuron, vol. 72, no. 4, pp. 665–678, Nov. 2011, doi: 10.1016/j.neuron.2011.09.006.

[6] K. J. Friston, “Functional and effective connectivity: a review,” Brain Connect, vol. 1, no. 1, pp. 13–36, 2011, doi: 10.1089/brain.2011.0008.

[7] M. Plitt, K. A. Barnes, and A. Martin, “Functional connectivity classification of autism identifies highly predictive brain features but falls short of biomarker standards,” Neuroimage Clin, vol. 7, pp. 359–366, 2015, doi: 10.1016/j.nicl.2014.12.013.

[8] R. Zebari, A. Abdulazeez, D. Zeebaree, D. Zebari, and J. Saeed, “A Comprehensive Review of Dimensionality Reduction Techniques for Feature Selection and Feature Extraction,” Journal of Applied Science and Technology Trends, vol. 1, no. 2, Art. no. 2, May 2020, doi: 10.38094/jastt1224.

[9] Y. Bengio, A. Courville, and P. Vincent, “Representation Learning: A Review and New Perspectives,” IEEE Transactions on Pattern Analysis and Machine Intelligence, vol. 35, no. 8, pp. 1798–1828, Aug. 2013, doi: 10.1109/TPAMI.2013.50.

[10] M. Dash and H. Liu, “Feature selection for classification,” Intelligent Data Analysis, vol. 1, no. 1, pp. 131–156, Jan. 1997, doi: 10.1016/S1088-467X(97)00008-5.

[11] J. Li et al., “Feature selection: A data perspective,” ACM Computing Surveys, vol. 50, no. 6, Dec. 2017, doi: 10.1145/3136625.

[12] C. Cameron et al., “The Neuro Bureau Preprocessing Initiative: open sharing of preprocessed neuroimaging data and derivatives,” Front. Neuroinform., vol. 7, 2013, doi: 10.3389/conf.fninf.2013.09.00041.

[13] J. Zhang, F. Feng, T. Han, X. Gong, and F. Duan, “Detection of Autism Spectrum Disorder using fMRI Functional Connectivity with Feature Selection and Deep Learning,” Cogn Comput, Jan. 2022, doi: 10.1007/s12559-021-09981-z.

[14] L. Shao, C. Fu, Y. You, and D. Fu, “Classification of ASD based on fMRI data with deep learning,” Cogn Neurodyn, vol. 15, no. 6, pp. 961–974, Dec. 2021, doi: 10.1007/s11571-021-09683-0.

[15] X. Bi, Y. Wang, Q. Shu, Q. Sun, and Q. Xu, “Classification of Autism Spectrum Disorder Using Random Support Vector Machine Cluster,” Frontiers in Genetics, vol. 9, 2018, Accessed: May 04, 2023. [Online]. Available: https://www.frontiersin.org/articles/10.3389/fgene.2018.00018

[16] X. Guo, K. C. Dominick, A. A. Minai, H. Li, C. A. Erickson, and L. J. Lu, “Diagnosing Autism Spectrum Disorder from Brain Resting-State Functional Connectivity Patterns Using a Deep Neural Network with a Novel Feature Selection Method,” Frontiers in Neuroscience, vol. 11, 2017, Accessed: May 04, 2023. [Online]. Available: https://www.frontiersin.org/articles/10.3389/fnins.2017.00460

[17] A. Kazeminejad and R. C. Sotero, “Topological Properties of Resting-State fMRI Functional Networks Improve Machine Learning-Based Autism Classification,” Frontiers in Neuroscience, vol. 12, 2019, Accessed: Mar. 18, 2023. [Online]. Available: https://www.frontiersin.org/articles/10.3389/fnins.2018.01018

[18] R. C. Sotero and J. M. Sanchez-Bornot, “Exploring Correlation-Based Brain Networks with Adaptive Signed Random Walks.” bioRxiv, p. 2023.04.27.538574, Apr. 27, 2023. doi: 10.1101/2023.04.27.538574.

[19] N. Tzourio-Mazoyer et al., “Automated anatomical labeling of activations in SPM using a macroscopic anatomical parcellation of the MNI MRI single-subject brain,” Neuroimage, vol. 15, no. 1, pp. 273–289, Jan. 2002, doi: 10.1006/nimg.2001.0978.

[20] C. Cameron et al., “Towards Automated Analysis of Connectomes: The Configurable Pipeline for the Analysis of Connectomes (C-PAC),” Front. Neuroinform., vol. 7, 2013, doi: 10.3389/conf.fninf.2013.09.00042.

[21] R. W. Cox, “AFNI: Software for Analysis and Visualization of Functional Magnetic Resonance Neuroimages,” Computers and Biomedical Research, vol. 29, no. 3, pp. 162–173, Jun. 1996, doi: 10.1006/cbmr.1996.0014.

[22] S. M. Smith et al., “Advances in functional and structural MR image analysis and implementation as FSL,” Neuroimage, vol. 23 Suppl 1, pp. S208–219, 2004, doi: 10.1016/j.neuroimage.2004.07.051.

[23] G. Grabner, A. L. Janke, M. M. Budge, D. Smith, J. Pruessner, and D. L. Collins, “Symmetric atlasing and model based segmentation: an application to the hippocampus in older adults,” Med Image Comput Comput Assist Interv, vol. 9, no. Pt 2, pp. 58–66, 2006, doi: 10.1007/11866763_8.

[24] J. Mazziotta et al., “A Four-Dimensional Probabilistic Atlas of the Human Brain,” Journal of the American Medical Informatics Association, vol. 8, no. 5, pp. 401–430, Sep. 2001, doi: 10.1136/jamia.2001.0080401.

[25] B. B. Avants, N. J. Tustison, G. Song, P. A. Cook, A. Klein, and J. C. Gee, “A reproducible evaluation of ANTs similarity metric performance in brain image registration,” NeuroImage, vol. 54, no. 3, pp. 2033–2044, Feb. 2011, doi: 10.1016/j.neuroimage.2010.09.025.

[26] Y. Behzadi, K. Restom, J. Liau, and T. T. Liu, “A component based noise correction method (CompCor) for BOLD and perfusion based fMRI,” Neuroimage, vol. 37, no. 1, pp. 90–101, Aug. 2007, doi: 10.1016/j.neuroimage.2007.04.042.

[27] C. E. Davey, D. B. Grayden, G. F. Egan, and L. A. Johnston, “Filtering induces correlation in fMRI resting state data,” Neuroimage, vol. 64, pp. 728–740, Jan. 2013, doi: 10.1016/j.neuroimage.2012.08.022.

[28] K. Murphy, R. M. Birn, D. A. Handwerker, T. B. Jones, and P. A. Bandettini, “The impact of global signal regression on resting state correlations: Are anti-correlated networks introduced?,” NeuroImage, vol. 44, no. 3, pp. 893–905, Feb. 2009, doi: 10.1016/j.neuroimage.2008.09.036.

[29] A. Di Martino et al., “The autism brain imaging data exchange: towards a large-scale evaluation of the intrinsic brain architecture in autism,” Mol Psychiatry, vol. 19, no. 6, pp. 659–667, Jun. 2014, doi: 10.1038/mp.2013.78.

[30] H. Wang et al., “Developmental brain structural atypicalities in autism: a voxel-based morphometry analysis,” Child Adolesc Psychiatry Ment Health, vol. 16, p. 7, Jan. 2022, doi: 10.1186/s13034-022-00443-4.

[31] D. Kim et al., “Overconnectivity of the right Heschl’s and inferior temporal gyrus correlates with symptom severity in preschoolers with autism spectrum disorder,” Autism Res, vol. 14, no. 11, pp. 2314–2329, Nov. 2021, doi: 10.1002/aur.2609.

[32] A. D. Bull, “Convergence Rates of Efficient Global Optimization Algorithms,” Journal of Machine Learning Research, vol. 12, no. 88, pp. 2879–2904, 2011.

[33] Y. LeCun, Y. Bengio, and G. Hinton, “Deep learning,” Nature, vol. 521, no. 7553, Art. no. 7553, May 2015, doi: 10.1038/nature14539.

[34] G. Dong and H. Liu, Feature Engineering for Machine Learning and Data Analytics. CRC Press, 2018.

[35] A. Holzinger, C. Biemann, C. S. Pattichis, and D. B. Kell, “What do we need to build explainable AI systems for the medical domain?” arXiv, Dec. 28, 2017. doi: 10.48550/arXiv.1712.09923.

[36] M. G. Preti, T. A. Bolton, and D. Van De Ville, “The dynamic functional connectome: State-of-the-art and perspectives,” NeuroImage, vol. 160, pp. 41–54, Oct. 2017, doi: 10.1016/j.neuroimage.2016.12.061.

[37] Z. S. Saad et al., “Trouble at Rest: How Correlation Patterns and Group Differences Become Distorted After Global Signal Regression,” Brain Connect, vol. 2, no. 1, pp. 25–32, Feb. 2012, doi: 10.1089/brain.2012.0080.

[38] A. Kazeminejad and R. C. Sotero, “The Importance of Anti-correlations in Graph Theory Based Classification of Autism Spectrum Disorder,” Frontiers in Neuroscience, vol. 14, 2020, Accessed: Sep. 09, 2022. [Online]. Available: https://www.frontiersin.org/articles/10.3389/fnins.2020.00676

[39] M. I. Al-Hiyali, N. Yahya, I. Faye, and A. F. Hussein, “Identification of Autism Subtypes Based on Wavelet Coherence of BOLD FMRI Signals Using Convolutional Neural Network,” Sensors, vol. 21, no. 16, Art. no. 16, Jan. 2021, doi: 10.3390/s21165256.

